# Requirement for PKC Epsilon in Kras-Driven Lung Tumorigenesis

**DOI:** 10.1101/2020.06.26.173690

**Authors:** Rachana Garg, Mariana Cooke, Shaofei Wang, Fernando Benavides, Martin C. Abba, Michelle Cicchini, David M. Feldser, Marcelo G. Kazanietz

## Abstract

Non-small cell lung cancer (NSCLC), the most frequent subtype of lung cancer, remains a highly lethal malignancy and one of the leading causes of cancer deaths worldwide. Mutant KRAS is the prevailing oncogenic driver of lung adenocarcinoma, the most common histological form of NSCLC. In this study, we examined the role of PKCε, an oncogenic kinase highly expressed in NSCLC and other cancers, in KRAS-driven tumorigenesis. Notably, database analysis revealed an association between PKCε expression and poor outcome in lung adenocarcinoma patients specifically having KRAS mutation. By generating a PKCε-deficient, conditionally activatable allele of oncogenic *Kras* (*LSL-Kras^G12D^*;PKCε^−/−^ mice) we were able to demonstrate the requirement of PKCε for *Kras*-driven lung tumorigenesis *in vivo*, which is consistent with the impaired transformed growth observed in PKCε-deficient KRAS-dependent NSCLC cells. Moreover, PKCε-knockout mice were found to be less susceptible to lung tumorigenesis induced by benzo[a]pyrene, a carcinogen that induces mutations in *Kras*. Mechanistic analysis using RNA-Seq revealed little overlapping for PKCε and KRAS in the control of genes/biological pathways relevant in NSCLC, suggesting that a permissive role of PKCε in *KRAS*-driven lung tumorigenesis may involve non-redundant mechanisms. Our results thus highlight the relevance and potential of targeting PKCε for lung cancer therapeutics.

## INTRODUCTION

Lung cancer is the leading cause of cancer-related deaths worldwide, accounting for 2.1 million new cases and 1.8 million deaths annually. Non-small cell lung cancer (NSCLC) is the most common form of lung cancer and constitutes ~85% of new diagnoses, with adenocarcinomas representing the predominant histological form. The disease is often detected at an advanced metastatic stage, resulting in poor patient prognosis (1). The vast majority of lung cancers are associated with long-term exposure to tobacco smoke and/or other environmental factors, such as benzo[a]pyrene (B[a]P) and other polycyclic aromatic hydrocarbons resulting from the combustion of organic matter. Most prevalent genetic alterations in lung adenocarcinomas include mutations in *KRAS* (~25%), *EGFR* (~15%), *PIK3CA*, *HER2* and *BRAF* (1-5%), *ALK* translocations (3-7%), and *MET* and *AXL* amplifications (1-5%). *KRAS* mutations, predominantly in codons 12-13, are found in approximately 1/3 of lung adenocarcinomas of smokers and have been associated with carcinogen exposure (1, 2). Targeted therapy against mutant *KRAS* has thus far been exceptionally challenging, thus stressing the crucial need to identify *KRAS* effectors as clinically actionable targets for disease management.

Protein kinase C epsilon (PKCε), a diacylglycerol (DAG) and phorbol ester regulated kinase, emerged as an oncogenic member of the PKC family, and has been widely regarded as a driver of cell survival, mitogenesis, motility and invasion. PKCε has been initially characterized as a transforming oncogene and subsequently recognized as a cancer biomarker, showing overexpression in multiple epithelial tumors, including lung cancer. Studies have shown that aberrantly expressed PKCε expression contributes to cancer initiation and progression (3–5). For example, prostate-specific PKCε overexpression in mice leads to the development of preneoplastic lesions that evolve to overt invasive adenocarcinoma in conjunction with loss of the tumor suppressor Pten (6). Transgenic PKCε overexpression in the mouse skin leads to the development of metastatic squamous cell carcinomas and enhances the susceptibility to UV radiation-induced skin cancer (7). The elevated PKCε levels in primary NSCLC tumors and cell lines has been linked to highly proliferative, survival and aggressive metastatic phenotypes (3–5,8). However, the association of PKCε expression with lung cancer oncogenic drivers or the susceptibility to carcinogenic insults remains unknown.

Here, we investigated the role of PKCε in lung cancer driven by mutant KRAS. Using a null mutant PKCε mouse model, we were able to define the involvement of PKCε in KRas-mediated and chemically-induced lung carcinogenesis, thus outlining a permissive role for this kinase in tumor development.

## MATERIALS AND METHODS

### Cell culture

Authenticated lung adenocarcinoma cell lines were obtained from ATCC, and cultured in RPMI medium supplemented with 10% FBS, 2 mM glutamine, 100 U/ml penicillin and 100 μg/ml streptomycin.

### RNAi, transfections and lentiviral infections

Cells were transfected with RNAi duplexes or infected with shRNA lentiviruses for either PKCε, KRAS or non-target control. For information about RNAi duplexes and lentiviruses, see Supplementary Data.

### Western blots and cell growth assays

Immunoblotting was carried out as described in (9). Details about the commercial antibodies are presented in Supplementary Data. Liquid colony formation (anchorage-dependent) and soft agar (anchorage-independent) growth assays have been previously described (6, see also Supplementary Data).

### Lung tumorigenesis studies

*LSL-Kras^G12D^* mice (The Jackson Laboratory) were crossed with PKCε KO mice (B6.129S4-*Prkce^tm1Msg/J^*, kindly provided by Dr. Robert Messing, University of Texas at Austin) to generate the different experimental genotypes. To initiate Kras^G12D^ transgene expression, Ad-Cre (1.2 × 10^4^ pfu) was inoculated intratracheally and lesion formation analyzed 32 week later.

For carcinogenesis studies, PKCε KO mice (B6.129S4-*Prkce^tm1Msg/J^*) introgressed into an A/J background were treated with B[a]P (1 mg/ml, *i.p.*, once/week for 4 weeks). Lesion analysis was done 20 weeks later. For speed congenic approach, see Supplementary Data.

### RNA-Seq

Directional RNA-Seq library construction and sequencing was carried out at the UPenn NGS Core. Detailed methodological and bioinformatics analysis is presented as Supplementary Data.

### Statistical analysis

ANOVA was performed using GraphPad Prism software built-in analysis tools. *p* values are indicated in the figure.

## RESULTS AND DISCUSSION

### *PRKCE* predicts poor outcome in mutant *KRAS* human lung adenocarcinoma

*KRAS* mutations are the most frequent oncogenic alterations reported in human lung adenocarcinoma and are often found in precancerous lesions such as atypical adenomatous hyperplasia (10). *Kras* mutations, primarily in codon 12, are also detected in spontaneous and chemically-induced mouse lung tumors (2). Since PKCε is up-regulated in human lung adenocarcinoma cell lines and tumors (4,5,8), we intended to ascertain if there exists any potential expression correlations with patient survival using publicly available datasets. For this analysis, we only selected samples with known *PRKCE* expression, *KRAS* mutational status and overall survival data, with a minimal follow-up of 10 months. Primary lung adenocarcinomas were first divided into low *PRKCE* and high *PRKCE* according to their median expression levels. Kaplan-Meier analysis of the overall population revealed no significant differences between patients with low and high *PRKCE* expression (Fig. 1A, *left panel*). Patients were then stratified according to their *KRAS* mutational status, totaling 352 wild-type *KRAS* and 73 mutant *KRAS* cases (43% *KRAS*^G12C^, 28% *KRAS*^G12V^, 10% *KRAS*^G12D^, 10% *KRAS*^G12A^, 5% *KRAS*^G12C^, 2% *KRAS*^G12R^ and 2% *KRAS*^G12Y^). Remarkably, whilst relatively similar outcomes were observed in patients with wild-type *KRAS* (Fig. 1A, *middle panel*), striking differences depending on *PRKCE* expression were found in patients with mutant *KRAS* tumors. Indeed, mutant *KRAS* patients with high *PRKCE* expression showed significantly worse survival compared to those having low *PRKCE* expression (~40% *vs.* 90% survival at ten years after diagnosis; Fig. 1A, *right panel*). Therefore, high *PRKCE* expression is indicative of poor prognosis in mutant *KRAS* patients.

**Figure 1.**
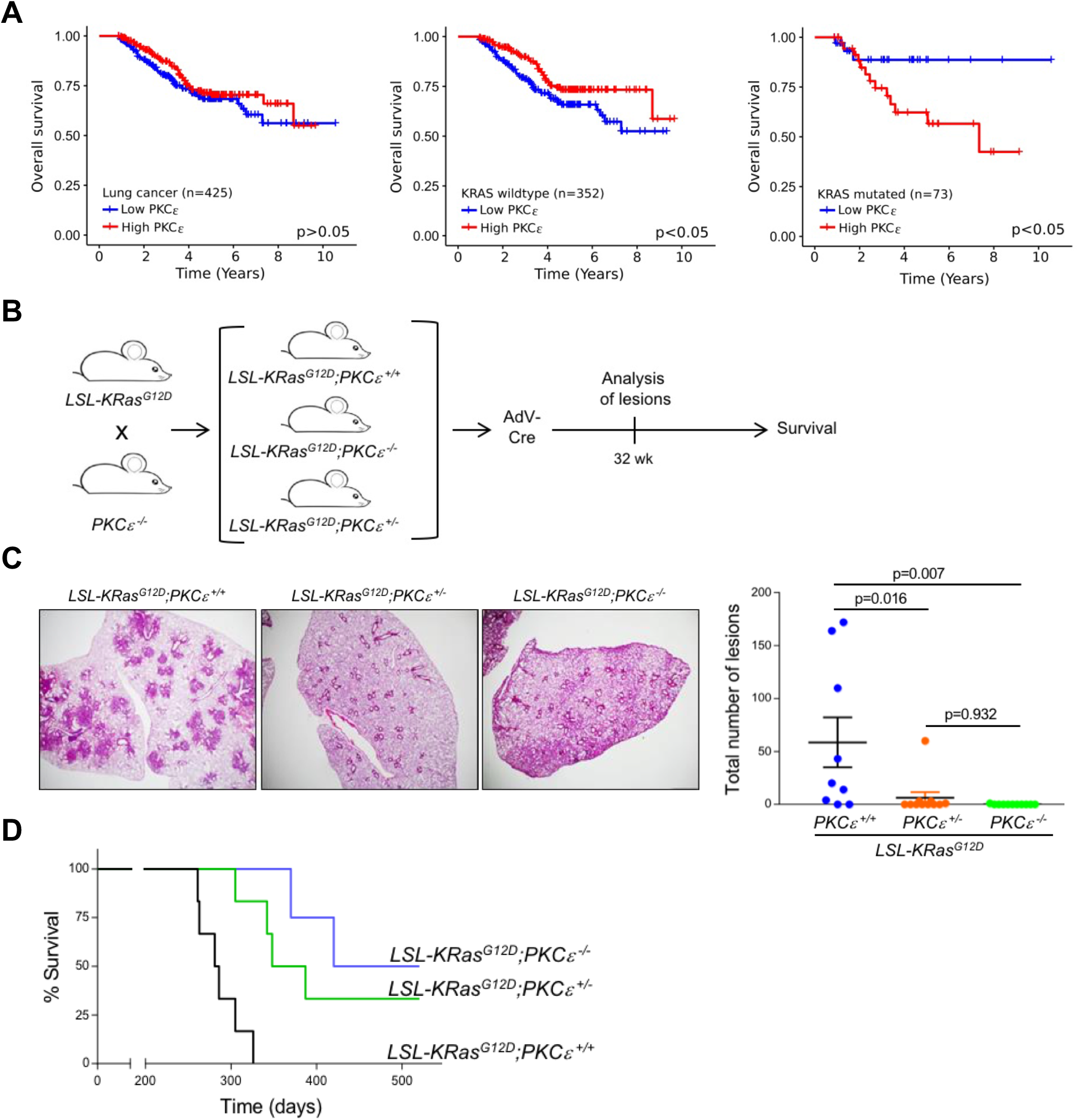
PKCε is required for the formation of Kras-driven lung lesions. *Panel A.* Kaplan-Meier analysis was carried out in a set of 425 patients with lung adenocarcinomas. Patients were categorized as “*PRKCE* high” (*red lines*) and “*PRKCE* low” (*blue lines*) according to the median expression of the Affymetrix 206248_at probe. Databases employed: GEO31210, E-MTAB923 and TCGA-LUAD. *Left panel*, all patients; *middle panel*, wild-type KRAS patients; *right panel*, mutant KRAS patients. *Panel B.* Schematic representation of the protocol for analysis of lung tumorigenesis and survival. To initiate Kras^G12D^ transgene expression, Ad-Cre (1.2 × 10^4^ pfu) was inoculated intratracheally in 6-8-week old mice of three different genotypes (*LSL-Kras^G12D^*;PKCε^+/+^, *LSL-Kras^G12D^*;PKCε^+/−^ and *LSL-Kras^G12D^*;PKCε^−/−^). *Panel C*. *Left panel*, representative photomicrographs of H&E stained lungs from *LSL-Kras^G12D^*;PKCε^+/+^, *LSL-Kras^G12D^*;PKCε^+/−^ and *LSL-Kras^G12D^*;PKCε^−/−^ mice. Magnification: 2X. *Right panel,* number of lesions in lungs from mice sacrificed 32 weeks after Ad-Cre delivery. Each point represents one mouse. Results are expressed as mean ± S.E.M. (n=8-11/group). *Panel D.* Kaplan-Meier analysis of mice survival for a period of 520 days post Ad-Cre delivery (n=8/group, p<0.05).

### Genetic deletion of PKCε inhibits *Kras*-mediated lung tumorigenesis in mice

While PKC isozymes, including PKCε, have been mostly implicated in tumor promotion, emerging evidence suggests a potential role for this kinase in tumor initiation (3, 6). Having observed an association between *PRKCE* expression and poor outcome in *KRAS* mutant patients, we next examined the involvement of PKCε in Kras-dependent formation of lung adenocarcinomas using a mouse model. Towards this goal, we employed a conditional mutant *Kras* mice (*LSL-Kras^G12D^*) (11) and examined if tumorigenesis ensues in a PKCε-null background. We therefore intercrossed *LSL-Kras^G12D^* mice with PKCε KO mice (B6.129S4-*Prkce^tm1Msg/J^*) to generate cohorts of *LSL-Kras^G12D^*;*Prkce*^+/+^, *LSL-Kras^G12D^*;*Prkce*^+/−^ and *LSL-Kras^G12D^*;*Prkce*^−/−^ mice. Expression of the oncogenic Kras^G12D^ allele in lungs was achieved following Ad-Cre intratracheal instillation of adenoviral Cre (Ad-Cre), which removes the Lox-Stop-Lox (LSL) cassette (Fig. 1B). *LSL-Kras^G12D^*;*Prkce*^+/+^ mice displayed characteristic lesions as reported previously in *Kras* mutant mice (11), including pulmonary adenoma and atypical adenomatous hyperplasia (46% of lesions) as well as bronchiolar/alveolar hyperplasia (54% of lesions). Remarkably, there was a significant reduction in the overall lesions in *LSL-Kras^G12D^*;*Prkce*^+/−^ mice, and essentially no lesions were detected in *LSL-Kras^G12D^*;*Prkce*^−/−^ mice (Figs. 1C). Thus, genetic deletion of the *Prkce* gene in mice impaired Kras-mediated lung tumorigenesis. Kaplan-Meier analysis revealed a median survival of 284 days for control *LSL-Kras^G12D^*;*Prkce*^+/+^ mice. However, loss of either one or two *Prkce* alleles significantly extended mice lifespan, with a median survival of 368 and 470 days, respectively (Fig. 1D).

### PKCε is required for B[a]P-induced lung carcinogenesis

Thus far, an unresolved question is whether PKCε contributes to lung tumorigenesis induced by chemical carcinogens. To answer this, we employed a known B[a]P-induced lung tumorigenesis model. Since B6.129S4 genetic background is poorly sensitive to the action of carcinogens, we crossed the PKCε allele into the A/J background, which is highly susceptible to chemical carcinogenesis (12). Using a speed congenic procedure, a 97% A/J background was achieved for the PKCε KO mice (Fig. S1).

Next, A/J PKCε^+/+^, PKCε^+/−^ and PKCε^−/−^ mice (6-8-weeks) were treated with B[a]P (Fig. 2A) and their lungs analyzed 20 weeks post carcinogen treatment (Fig. 2B). Nearly all PKCε^+/+^ mice (95%) developed lung lesions. Remarkably, the incidence of lesions was markedly diminished upon loss of one or two *Prkce* alleles. The incidence of B[a]P-induced adenomas in PKCε^+/+^, PKCε^+/−^ and PKCε^−/−^ mice was 76%, 50% and 31%, respectively. A similar trend was observed for the incidence of hyperplastic lesions developed in the various groups (Fig. 2C). An average of 4.5 lung lesions was found in PKCε^+/+^ mice, whereas the lesion multiplicity was significantly reduced in PKCε^+/−^ and PKCε^−/−^ mice (Fig. 2D). The observed inhibition of chemically-induced lung tumor formation even upon the loss of a single *Prkce* allele is a strong indicator that PKCε expression levels are key to determine susceptibility to the chemical carcinogen.

**Figure 2.**
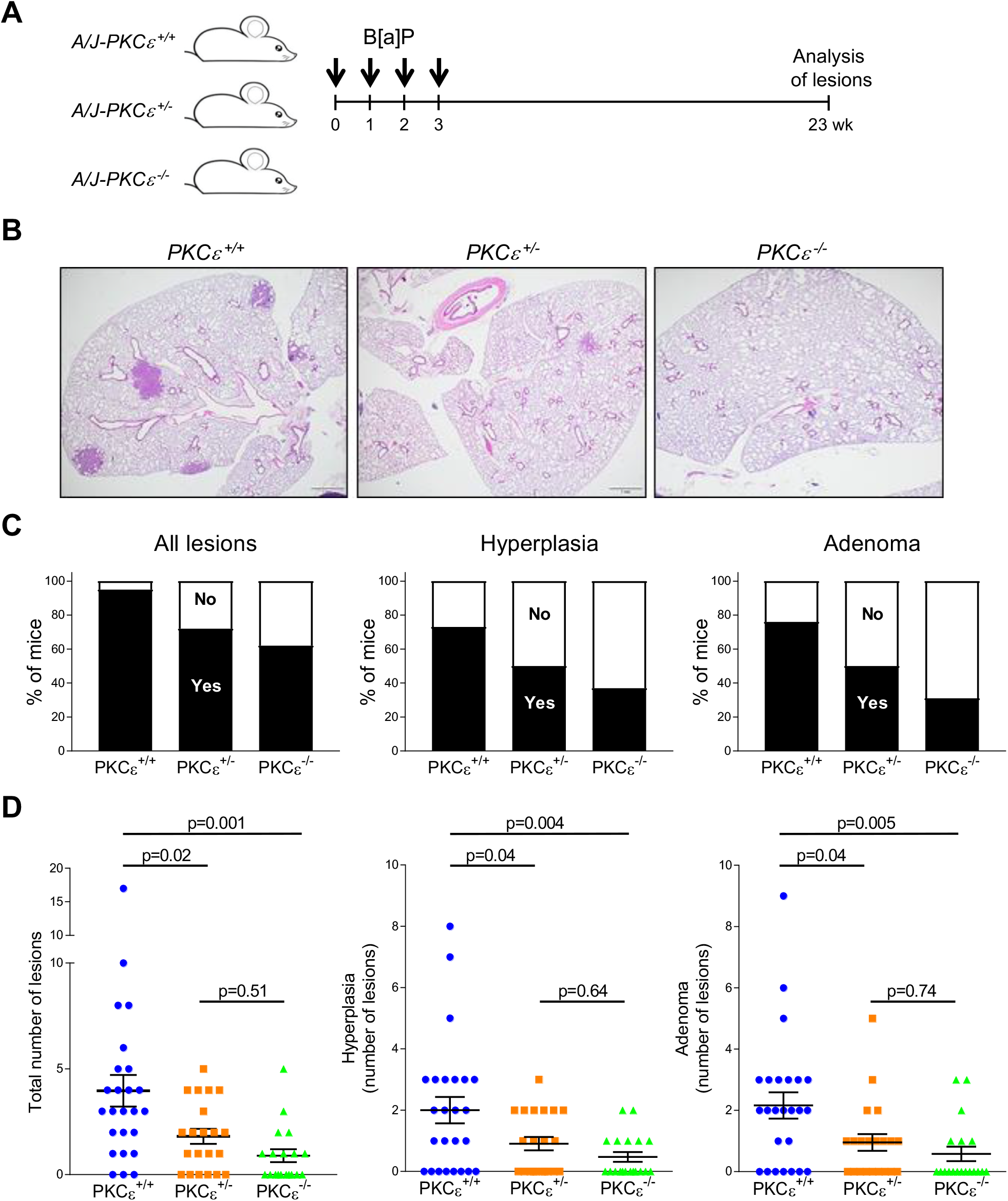
B[a]P-induced lung tumorigenesis is impaired in PKCε KO mice. PKCε KO mice (B6.129S4-*Prkce^tm1Msg/J^*) were introgressed into an A/J background using a speed congenic procedure (see Fig. S1). PKCε^+/+^, PKCε^+/−^ and PKCε^−/−^ mice (6-8-week old) in A/J background were treated with B[a]P (1 mg/ml, *i.p.*, once/week for 4 weeks) and sacrificed 20 weeks later. Excised lungs were subjected to H&E staining and histopathological analysis at the University of Pennsylvania Comparative Pathology Core. *Panel A.* Scheme depicting the experimental approach. *Panel B.* Representative photomicrographs of H&E stained lungs from PKCε^+/+^, PKCε^+/−^ and PKCε^−/−^ mice 20 weeks after B[a]P treatment. Magnification: 2X. *Panel C.* Incidence of total lesions, hyperplastic lesions and adenomas. *Panel D.* Analysis of multiplicity of lesions in different groups. Each point represents one mouse. Results are expressed as mean ±S.E.M. (n=16-25/group).

### Silencing PKCε expression reduces features of transformation of *KRAS* mutant NSCLC cells

To further assess the involvement of PKCε in *KRAS*-mediated phenotypes, we examined the PKCε requirement for growth of *KRAS* mutant human lung adenocarcinoma cells. Four different siRNAs were used to silence the expression of either KRAS (K1-K4) or PKCε (ε1-ε4 in H2009 cells (Fig. 3A, *left panel*). RNAi duplexes that cause a major knockdown in KRAS expression (K1 and K3) led to a prominent reduction in anchorage-dependent growth relative to cells transfected with a non-target control (NTC), as expected from the KRAS dependency of this cell line (13). A smaller effect was observed with RNAi duplexes that were less efficient to silence KRAS (K2 and K4). Similar decrease in H2009 cells anchorage-dependent growth was observed upon PKCε depletion, albeit the extent varied with different RNAi duplexes (Fig. 3A, *right panel*). Inhibition of colony formation following KRAS or PKCε RNA silencing was also observed in KRAS mutant NSCLC cell lines H358 and H441 (Figs. S2A, S2B, S2D and S2E). Notably, analysis of anchorage-independent growth in soft agar also revealed a consistent dependency on both KRAS and PKCε in H2009 (Fig. 3B) and H358 cells (Fig. S2C). H441 cells did not form colonies in soft agar (data not shown). For validation, we generated five PKCε-stably depleted H2009 cell lines using shRNA lentiviruses. These cell lines displayed in all cases reduced anchorage-dependent and anchorage-independent growth relative to cells transduced with NTC shRNA lentivirus (Fig. S3). Taken together, our data strongly support the contention that PKCε is required for KRAS-dependent transformed growth of NSCLC cells.

**Figure 3.**
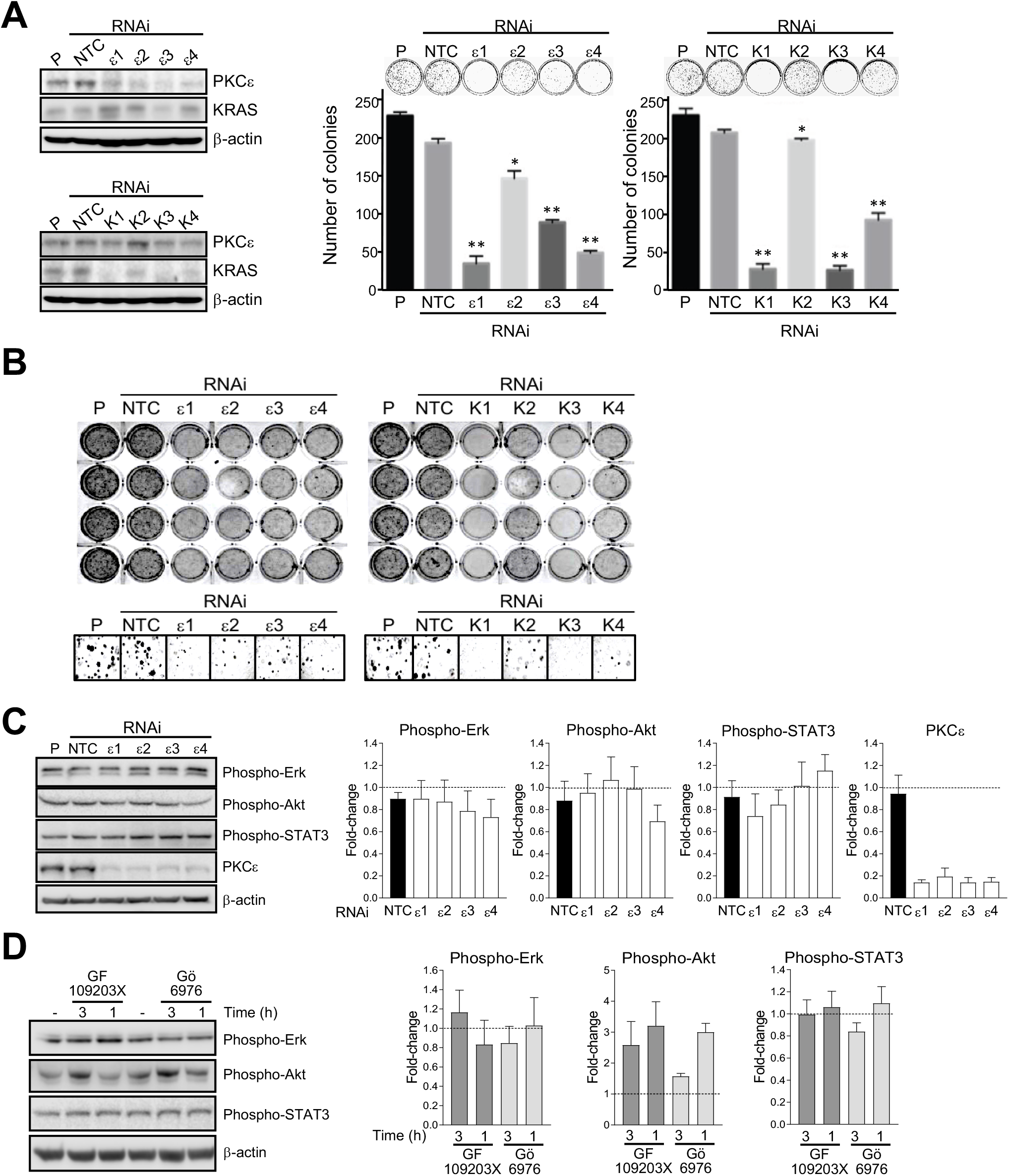
PKCε is required for clonogenic and transformed growth of KRAS-mutant NSCLC cells. *Panel A.* Expression of PKCε and KRAS in H2009 cells subjected to RNAi, as determined by Western blot (*left panel*). Anchorage-dependent growth was determined by means of a liquid colony formation. A representative experiment and quantitative analysis are shown. Results are expressed as mean ± S.E.M. (n=4). *, p<0.05 *vs.* NTC; **, p<0.01 *vs.* NTC (*right panel*). *Panel B.* Anchorage-independent growth in soft agar. Representative pictures of plates (*upper panels*) and photomicrographs of colonies *(lower panels*) are shown. Similar results were observed in two additional experiments. *Panel C.* H2009 cells were subjected to PKCε RNAi with duplexes ε1, ε2, ε3 and ε4. Forty-eight h later, cells were lysed and subject to Western blot analysis with the indicated antibodies. *Left panel*, representative experiment. *Right panels*, densitometric analysis of 3 independent experiments. Results are expressed as mean ± S.E.M. *Panel D.* H2009 cells were treated with PKC inhibitors GF109203X (3 μM) and Gö6976 (3 μM) for either 1 or 3 h. *Left panel*, representative Western blots with the indicated antibodies. *Right panels*, densitometric analysis of 3 independent experiments. Results were expressed as mean ± S.E.M.

Studies implicated PKCs in mitogenic and survival signaling in cancer cells (3–5). However, whether PKCε mediates signaling events in KRAS mutant NSCLC cells remains undetermined. To our surprise, silencing PKCε in KRAS mutant H2009 cells failed to reduce phosphorylated (active) levels of Erk, Akt and STAT3, well-established downstream effectors of KRAS (Fig. 3C). Likewise, the activation status of these effectors was not reduced upon treatment with either the “pan” PKC inhibitor (GF109203X) or the cPKC inhibitor Gö6976 (Fig. 3D). Rather, phospho-Akt levels were slightly elevated upon PKC inhibitors treatment. This effect is consistent with the known inhibition of Akt by PKCα(14), the only cPKC expressed in these cells (9). Similar results were observed in H358 and H441 cells (data not shown). Therefore, regardless of the PKCε requirement for NSCLC cell growth and lung tumorigenesis, this kinase or other PKCs may not act as a downstream effector of KRAS for key mitogenic and survival signaling pathways.

### Differential control of gene expression by PKCε and KRAS in NSCLC cells

As a next step in our search for potential common mechanisms by which KRAS and PKCε regulate transformed growth in NSCLC cells, we explored global gene expression changes. While KRAS-dependent signatures have been established, including in lung cancer (15, 16), studies suggested a limited involvement of PKCε in controlling gene expression compared to other PKC family members (9, 17). Using RNA-Seq, we carried out gene expression analysis in H2009 cells subjected to PKCε or KRAS depletion with two different duplexes in each case. To identify differentially regulated expression, we used the edgeR test (FDR<0.05, cut off: 2-fold change relative to NTC). This analysis revealed 260 differentially regulated genes (148 up- and 112 down-regulated) by both KRAS RNAi duplexes, whereas relatively less effect was observed upon PKCε silencing (73 deregulated genes, 38 up- and 35 down-regulated). As expected, KRAS and PKCε levels were among the down-regulated genes, with nearly complete depletion with each RNAi duplex (Fig. 4A). Heatmaps of deregulated transcripts are shown in Figs. 4B, and a complete list of genes is presented in Table S1. The limited involvement of PKCε in gene expression agrees with our recent study in NSCLC cells using the PKCε-specific ligand AJH-836 (9). Interestingly, a small overlap (15 genes) was found between KRAS- and PKCε-regulated genes, in all cases genes up-regulated upon specific silencing (Fig. 4C).

**Figure 4.**
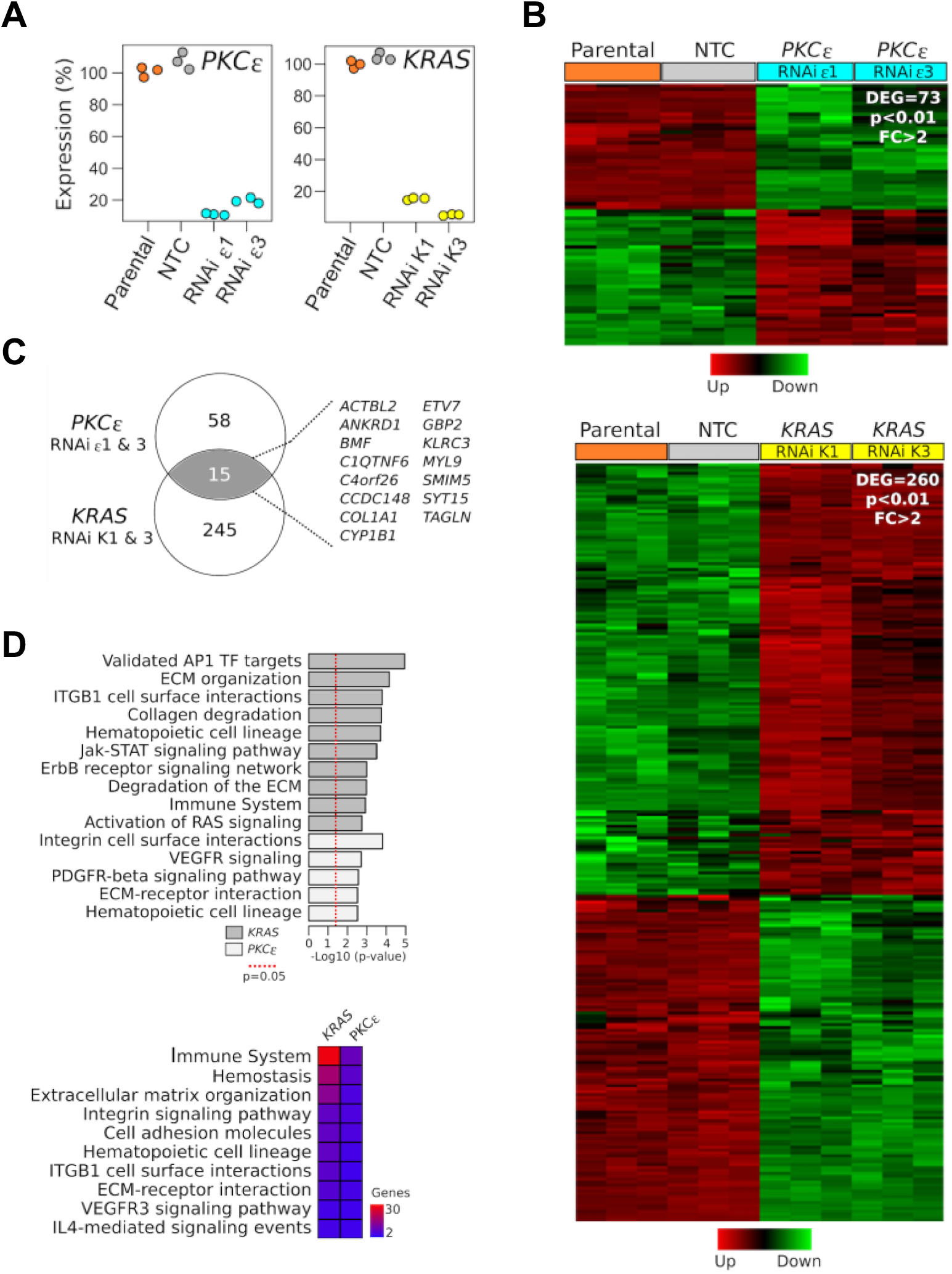
Gene expression analysis in PKCε- and KRAS-depleted NSCLC cells. H2009 cells were subjected to PKCε RNAi (ε1 or ε3 duplexes) or KRAS RNAi (K1 or K3 duplexes), and RNA-Seq analysis for gene expression was performed 48 h after transfection. As controls we used NTC RNAi and parental cells. Three replicates were done for each condition. *Panel A.* Validation of PKCε and KRAS silencing from the RNA-Seq data in individual samples. *Panel B.* Heatmap of 73 deregulated genes (*DEG*) in H2009 cells subjected to PKCε RNAi depletion (*upper panel*) or KRAS RNAi depletion (*lower panel*). The color scale at the bottom of the heatmap is used to represent expression level (*green*, low expression; *red*, high expression). Fold-change > 2; FDR < 0.01). *Panel C.* Venn diagram of transcripts commonly modulated among PKCε and KRAS silenced H2009 cells. *Panel D.* Top bioprocess enriched in the PKCε and KRAS gene expression signatures. The red dotted line indicates the cut off for statistical significance (p<0.05) (*upper panel*). A comparative analysis of the bioprocesses commonly enriched across the PKCε and KRAS regulated genes was done using the InnateDB resource (*lower panel*).

Using InnateDB, we carried out automated annotation and functional enrichment analysis of the differentially expressed genes. The top statistically significant bioprocesses regulated by KRAS included AP1 transcription factor targets, extracellular matrix (ECM) organization and degradation, integrin-β1 cell surface interaction, as well as Jak-STAT, ErbB and RAS signaling pathways. Despite the small number of genes affected by PKCε silencing, we identified a few statistically significant bioprocesses regulated by this kinase, including integrin cell surface/ECM interactions and VEGFR/PDGFR signaling pathways (Fig. 4D, *upper panel*; see Table S2 for complete list). A few bioprocesses were found to be commonly regulated by KRAS and PKCε, though the number of genes involved is relatively small, particularly in the case of PKCε (Fig. 4D, *lower panel*). Therefore, it is reasonable to speculate that the requirement of PKCε for KRAS-mediated growth and tumorigenesis does not rely on common transcriptional genetic programs.

### Final remarks

Our findings revealed a functional association between KRAS and PKCε contributing to the development of lung cancer. The reduced number of Kras-driven and B[a]P-induced lung lesions in a PKCε-deficient background strongly argues for a key permissive role of this kinase in tumor initiation. Although a comprehensive mechanistic assessment of the hierarchical relationship between KRAS and PKCε has yet to be pursued, the fact that loss of PKCε does not substantially affect the activation status of KRAS signaling effectors or the expression of KRAS-regulated genes argues against PKCε acting as a KRAS downstream effector. Rather, we speculate that both KRAS and PKCε act in a coordinated manner through parallel pathways, whereby PKCε likely provides a facilitating input (*i.e.* a survival signal) for oncogenesis. The overlap in KRAS and PKCε bioprocesses, including those related to ECM, integrins and adhesion, together with the known PKCε involvement in integrin function and anoikis resistance (3,18,19), may throw light on the molecular basis of the KRAS-PKCε functional interaction.

NSCLC tumors display elevated PKCε levels (3–5,8) According to our dataset analysis, PKCε may represent a prognostic biomarker of poor outcome specifically in KRAS mutant lung adenocarcinoma patients. We speculate that PKCε contributes to disease development in a coordinated manner with specific genetic alterations, as recently shown for Pten-deficient prostate cancers (6), and serves as a marker for stratification of patients based on its expression. Notably, the reported reversion of Ras-mediated transformed phenotypes by pharmacological inhibition of PKCε (20) highlights unique opportunities for therapeutic interventions involving PKCε targeting. Given the relevance of KRAS oncogenic signaling in the progression of lung adenocarcinomas, digging into the intricacies of KRAS/PKCε interactions may uncover novel cross-talks between these players and help rationalize the potential for developing PKCε targeted therapies for oncogenic Ras-driven malignancies.

