## Supplemental File for "Requirement for PKC Epsilon in Kras-Driven Lung Tumorigenesis"

**Figure S1**

| Stage | A/J allele (%) |
|---|---|
| N1 | 0 |
| N2 | 70-81 |
| N3 | 91-95 |
| N4 | 97 |

**Figure S1. Generation of PKCε knockout mice in A/J background.** The *Prkce*$^{tm1Msg}$ null allele was transferred from a mixed B6;129S4 background onto A/J inbred background by marker-assisted backcrossing (speed congenics). The table shows the percentage of A/J allele of each backcross generation.

**Figure S2**

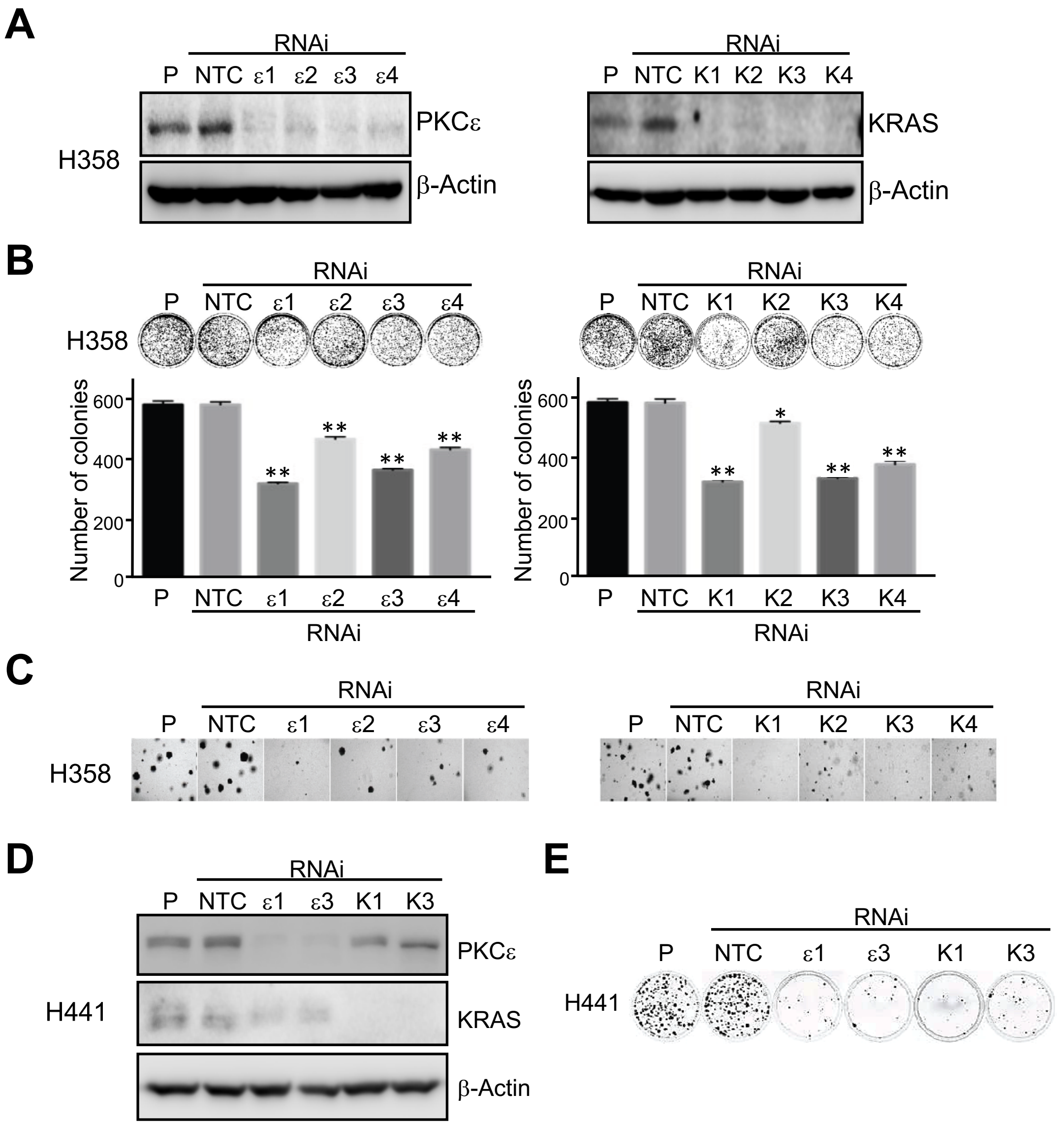


**Figure S2. PKCε- and KRAS-dependency for clonogenic and transformed growth in H358 and H441 KRAS-mutant NSCLC cells.** H358 and H441 lung adenocarcinoma cells were transfected with the RNAi duplexes for either PKCε or KRAS indicated in the figure. *Panel A.* Expression of PKCε and KRAS in H358 cells subjected to PKCε RNAi (*left panel*) or KRAS RNAi (*right panel*). *Panel B.* For the liquid colony formation assays, $3 \times 10^3$ H358 cells were plated in 100 mm plates. Colonies were determined 10 days later after staining with 0.7% methylene blue in 50% ethanol. Results are expressed as mean ± S.E. (n=4). *, $p<0.05$ *vs.* NTC; **, $p<0.01$ *vs.* NTC. *Panel C.* Representative photomicrographs of colonies in H358 anchorage-independent growth assays in soft agar. *Panel D.* Expression of PKCε and KRAS in H441 cells subjected to PKCε or KRAS RNAi, as determined by Western blot. *Panel E.* A representative liquid colony formation assay in H441 cells is shown. Anchorage-independent growth could not be evaluated, since H441 cells do not form colonies in soft agar.

**Figure S3**

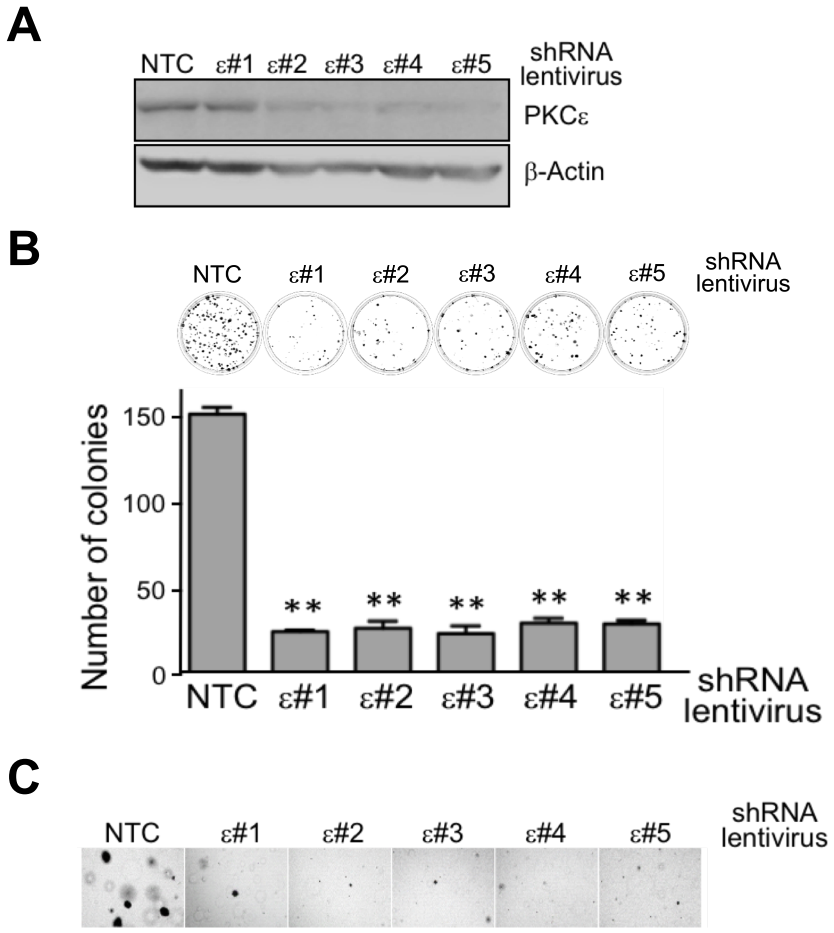


**Figure S3. Effect of stable PKCε depletion on anchorage-dependent and anchorage-independent H2009 cell growth.** H2009 cells were infected with shRNA lentiviruses for PKCε (*#1*-*#5*) or non-target control (*NTC*) lentivirus, and stable cell lines were selected with puromycin. *Panel A*. Western blot analysis for PKCε and KRAS in the stable cell lines. *Panel B*. Liquid colony formation assays for anchorage-dependent growth. For these assays, 500 cells were plated in 100 mm plates. Medium was replaced twice a week, and after 12 days colonies were stained with 0.7% methylene blue in 50% ethanol. *Upper panel*, representative experiment; *lower panel*, quantification of colonies. Results are expressed as mean ± S.E. (n=4). **p<0.01 *vs.* NTC. *Panel C*. Soft agar colony formation assays for anchorage-independent growth. Cells ($3 \times 10^3$) were plated in 24-well dishes in 0.35% agar over a 0.5% agar layer. After 10 days, the plates were stained with MTS. Representative photomicrographs of colonies are shown.

**SUPPLEMENTARY DATA**

### REQUIREMENT FOR PKC EPSILON IN KRAS-DRIVEN LUNG TUMORIGENESIS

Rachana Garg, Mariana Cooke, Shaofei Wang, Fernando Benavides, Martin C. Abba, Michelle Cicchini, David M. Feldser and Marcelo G. Kazanietz

#### SUPPLEMENTAL TABLE LEGENDS

**Table S1.** Expression profiles for genes regulated by silencing either PKCε or KRAS, grouped into either up- or down-regulated genes.

**Table S2.** Bioprocess enriched in the PKCε and KRAS gene expression signatures.

#### SUPPLEMENTAL EXPERIMENTAL PROCEDURES

##### Antibodies used for Western blot

The following antibodies and dilutions were used: anti-PKCε (1:1,000, Cell Signaling Technology, Danvers, MA), anti-KRAS (sc-30, 1:1,000, Santa Cruz Biotechnology, Dallas, TX), anti-phospho-Akt, anti-phospho-ERK, anti-phospho-STAT3, anti-PKCε (1:1:000, Cell Signaling Technology) and anti-β-actin (1:10,000, Sigma-Aldrich, St. Louis, MO). Densitometric analysis was done with an Odyssey Fc system (LI-COR Biotechnology, Lincoln, NE).

##### RNAi, transfections and lentiviral infections

Cells were transfected with RNAi duplexes for either PKCε or KRAS from Dharmacon (Lafayette, CO) using Lipofectamine RNAi Max (Invitrogen, Carlsbad, CA). RNAi's for PKCε were as follows: ON-TARGETplus PRKCE siRNA J-004653-06 (*ε1*), J-004653-07 (*ε2*), J-004653-08 (*ε3*) and J-004653-09 (*ε4*). For KRAS, RNAi's were as follows: ON-TARGETplus KRAS siRNA J-005069-08 (*K1*), J-005069-09 (*K2*), J-005069-10 (*K3*) and J-005069-11 (*K4*). As a non-target control (*NTC*), we used D-001810-02-05 ON-TARGETplus non-targeting siRNA.

For some experiments, H2009 cells were infected with shRNA lentiviruses for PKCε (MISSION shRNA Lentiviral Transduction particles, Sigma). Clone IDs are as follows: TRCN0000195553 (*#1*), TRCN0000000845 (*#2)*, TRCN0000000846 (*#3*), TRCN0000000847 (*#4*) and TRCN0000000848 (*#5*). As non-target control (*NTC*), we used MISSION pLKO.1-puro Control Transduction Particles (Catalog # SHC001V). Upon infection, stable cell lines were selected with puromycin (1 μg/ml).

#### Cell growth assays

For anchorage-dependent growth, 500 cells were plated in 100 mm plate for assessing liquid colony formation. Medium was replaced twice a week, and after 10 days colonies were stained with 0.7% methylene blue in 50% ethanol. After washing out the dye, colonies containing > 50 cells were counted. The colony formation efficiency was determined as the ratio of the colony number to the number of cells seeded. For the analysis of anchorage-independent growth, $3 \times 10^3$ cells were plated in 24-well dishes in 0.35% agar over a 0.5% agar layer. After 10 days, the plates were stained with 3-(4,5-dimethylthiazol-2-yl)-5-(3-carboxymethoxyphenyl)-2-(4-sulfophenyl)-2H-tetrazolium (MTS).

#### Speed congenic approach to generate A/J PKCε knockout mice.

The *Prkce*$^{tm1Msg}$ null allele (targeted using 129S4/SvJae derived RF8 ES cells) was transferred from a mixed B6;129S4 background onto A/J inbred background (Stock No: 000646) by marker-assisted backcrossing (speed congenics). Briefly, after the initial round of intercross and backcross (donor female), N2 males (heterozygous carriers for the *Prkce* null allele) were genotyped with 80 polymorphic microsatellite markers distributed throughout the genome in order to select the "best" males (*i.e.*, the ones that harbor the least amount of donor mixed genome). From this point on, we backcrossed the "best" male of each backcross generation (N2, N3, N4) with pure A/J females to produce the congenic strain A/J.B6-*Prkce*$^{tm1Msg}$ with 96% A/J background. Microsatellite genotyping was carried out in the Laboratory Animal Genetic Services located at MD Anderson Cancer Center, Smithville (TX).

#### RNA-Seq and bioinformatics

H2009 cells were subjected to PKCε RNAi (ε1 or ε3 duplexes) or KRAS RNAi (K1 or K3 duplexes). As controls we used NTC RNAi and parental cells. Three replicates were done for each condition. Forty-eight h later, total RNA was isolated from subconfluent plates using the RNeasy kit (Qiagen, Valencia, CA). RNA concentration and integrity were determined on an Agilent 2100 Bioanalyzer (Agilent Technologies). Samples were processed for directional RNA-Seq library construction and sequencing at the NGS Core facility of the Perelman School of Medicine, University of Pennsylvania. A 100 nt singled-end sequencing was performed using an Illumina HiSeq 4000 platform, and ~20 million reads per sample were obtained. The short-sequenced reads were mapped to the human reference genome (hg19) with the RNA-Seq unified mapper (RUM) V2.0.5. 06 (http://cbil.upenn.edu/RUM/). The R/Bioconductor packages were employed to accurately calculate the gene expression abundance using the aligned BAM files.

To identify differentially expressed genes (log2 fold change > ±1, false discovery rate <0.05) between various treatment groups and vehicle samples we employed the edgeR Bioconductor package based on the normalized log2 based count per million values. Raw data has been submitted to the NCBI GEO database. Comparative analysis of the bioprocesses commonly enriched across the PKCε and KRAS regulated genes was done using the InnateDB resource.

### SUPPLEMENTAL FIGURE LEGENDS

**Figure S1. Generation of PKCε knockout mice in A/J background.** The *Prkce*$^{tm1Msg}$ null allele was transferred from a mixed B6;129S4 background onto A/J inbred background by marker-assisted backcrossing (speed congenics). The table shows the percentage of A/J allele of each backcross generation.

**Figure S2. PKCε- and KRAS-dependency for clonogenic and transformed growth in H358 and H441 KRAS-mutant NSCLC cells.** H358 and H441 lung adenocarcinoma cells were transfected with the RNAi duplexes for either PKCε or KRAS indicated in the figure. *Panel A.* Expression of PKCε and KRAS in H358 cells subjected to PKCε RNAi (*left panel*) or KRAS RNAi (*right panel*). *Panel B.* For the liquid colony formation assays, $3 \times 10^3$ H358 cells were plated in 100 mm plates. Colonies were determined 10 days later after staining with 0.7% methylene blue in 50% ethanol. Results are expressed as mean ± S.E. (n=4). *, $p<0.05$ *vs.* NTC; **, $p<0.01$ *vs.* NTC. *Panel C.* Representative photomicrographs of colonies in H358 anchorage-independent growth assays in soft agar. *Panel D.* Expression of PKCε and KRAS in H441 cells subjected to PKCε or KRAS RNAi, as determined by Western blot. *Panel E.* A representative liquid colony formation assay in H441 cells is shown. Anchorage-independent growth could not be evaluated, since H441 cells do not form colonies in soft agar.

**Figure S3. Effect of stable PKCε depletion on anchorage-dependent and anchorage-independent H2009 cell growth.** H2009 cells were infected with shRNA lentiviruses for PKCε (*#1-#5*) or non-target control (*NTC*) lentivirus, and stable cell lines were selected with puromycin. *Panel A*. Western blot analysis for PKCε and KRAS in the stable cell lines. *Panel B*. Liquid colony formation assays for anchorage-dependent growth. For these assays, 500 cells were plated in 100 mm plates. Medium was replaced twice a week, and after 12 days colonies were stained with 0.7% methylene blue in 50% ethanol. *Upper panel*, representative experiment; *lower panel*, quantification of colonies. Results are expressed as mean ± S.E. (n=4). **$p<0.01$ *vs.* NTC. *Panel C*. Soft agar colony formation assays for anchorage-independent growth. Cells ($3 \times 10^3$) were plated in 24-well dishes in 0.35% agar over a 0.5% agar layer. After 10 days, the plates were stained with MTS. Representative photomicrographs of colonies are shown.
